# Reconstruction and exploitation of a dedicated Genome-Scale Metabolic Model of the human pathogen *C. neoformans*

**DOI:** 10.1101/2025.04.02.646762

**Authors:** Romeu Viana, Diogo Couceiro, William Newton, Luís Coutinho, Oscar Dias, Carolina Coelho, Miguel Cacho Teixeira

## Abstract

*C. neoformans* is notorious for causing severe pulmonary and central nervous system infections, particularly in immunocompromised patients. High mortality rates, associated with its tropism and adaptation to the brain microenvironment and its drug resistance profile, makes this pathogen a public health threat and a World Health Organization (WHO) priority.

In this study, we reconstructed GSMM iRV890 for *C. neoformans var. grubii*, providing a promising platform for the comprehensive understanding of the unique metabolic features of *C. neoformans*, and subsequently shedding light on its complex tropism for the brain microenvironment and potentially informing the discovery of new drug targets. The GSMM iRV890 model is openly available in the SBML format, and underwent validation using experimental data for nitrogen and carbon assimilation, as well as specific growth and glucose consumption rates. Based on the comparison with GSMMs available for other pathogenic yeasts, unique metabolic features were predicted for *C. neoformans*, including key pathways shaping the dynamics between *C. neoformans* and the human host, and underlying its adaptation to the brain environment. Finally, predicted essential genes from the validated model are explored herein as potential novel antifungal drug targets.

## 1. Introduction

Cryptococcal meningitis is a disease caused by a few pathogenic basidiomycetous yeast species, namely *Cryptococcus neoformans* (*C. neoformans)* and *Cryptococcus gatii*. Cryptococcosis is caused by three *Cryptococcus* species/variants, *C. neoformans var. grubii* (serotype A), responsible for 95% of *Cryptococcus* infections worldwide [1]; *C. neoformans var. neoformans* (serotype D) and *Cryptococcus gattii* (serotypes B and C) geographically restricted to tropical and/or subtropical regions [2].

These species are notorious for inducing severe pulmonary and central nervous system infections [3]. While these pathogens are harmless in healthy individuals, they poses a serious threat to immunocompromised patients, especially those with acquired immunodeficiency syndrome (AIDS) or those undergoing immunosuppressive therapies, causing severe meningoencephalitis and other serious neurological complications [4–6]. The latest systematic review, using data from more than 120 countries, estimates that cryptococcal meningitis affects 190 000 people worldwide annually, being associated with a mortality rate of 76% [7]. Cryptococcal infections are commonly treated with combination therapy, usually flucytosine in combination with amphotericin B in a first induction stage, followed by consolidation and long-term maintenance with high dose fluconazole [8]. Anti-cryptococcal monotherapy is not considered optimal, as it carries the risk of drug resistance [9]. Still, fluconazole monotherapy is still used, mostly due to limited drug access. An increase in fluconazole resistance among *C. neoformans* isolates was observed in past decades [10,11]. Fluconazole resistance is particularly notorious in isolates from relapse disease [12]. Despite verified *in vitro* susceptibility, echinocandins are not used clinically to treat cryptococcosis due to intrinsic resistance *in vivo* [13], attributed to echinocandins inability to penetrate the blood-brain barrier. Another possible contributor to echinocandin resistance in *Cryptococcus* species is fungal cell wall melanization, through the action of a fungal laccase, which uses the L-DOPA and dopamine found in the human brain as precursors [14]. Melanin is an important virulence factor in *C. neoformans* since it can neutralize oxidative stress radicals [15], as well as some toxic compounds, including some antifungal drugs, such as caspofungin and amphotericin B [16,17].

*C. neoformans* is widely spread in the environment, with worldwide distribution, in bird guano, soils and trees. Fungal particles are then inhaled by humans and other mammals [2]. This pathogen is characterized by their high resistance to harsh environments in nature and in mammalian hosts [18], and after inhalation into the host’s lungs, *Cryptococccus* can stay in a dormant latent granulomatous form for long periods of time [3]. However, tropism for the central nervous system is not yet fully understood [2,3]. Despite being a public health threat and a WHO priority pathogen [19], *C. neoformans* still has many aspects of its peculiar metabolism associated with the central nervous system and interactions with the host that remain poorly understood [20].

In this work, iRV890 the first reconstructed GSMM for the human pathogen *C. neoformans var. grubii*, a frequent variant of these pathogenic species, is presented. To facilitate usage by other researchers, the model is provided in the widely used SBML format. Model validation was conducted using experimental data for nitrogen and carbon assimilation from phenotypic arrays covering 222 different sources [21]. Specific growth and glucose consumption rates were experimentally determined in order to quantitatively validate the model’s predictive power. A set of essential genes derived from the validated model is predicted and discussed in terms of their potential as novel antifungal drug targets. An additional comparison, with GSMM’s for other pathogenic yeast species and *S. cerevisiae* was performed regarding the gene essentiality prediction and unique metabolic features of *C. neoformans*. Some peculiar characteristics and pathways of this fungus relevant to its pathogenicity are also discussed based on our findings. The iRV890 model provides a promising platform for global elucidation of the metabolic features of *C. neoformans var. grubii*, with expected impact in guiding the identification of new drug targets and understanding the complex metabolism of this pathogen in the context of the human brain.

## 2. Materials and Methods

### 2.1. Model Development

The genome-scale metabolic model of *C. neoformans var. grubii H99*, designated as iRV890, was reconstructed using *merlin* 4.0.5 [22] following the methodology described elsewhere [23] and OptFlux 3.0 [24], for curation and subsequent validation stages. All computational analyses were executed utilizing the IBM CPLEX 12.10 solver.

### 2.2. Genome Annotation and Assembling of the Metabolic Network

The genome sequence of the *C. neoformans var. grubii* was retrieved from NCBI’s Assembly database, with the accession number ASM1180120v1 [25] and the Taxonomy ID 235443 from NCBÍs Taxonomy database [26]. The genome-wide functional annotation was based on taxonomy and frequency of similar sequences through remote DIAMOND alignment [27] and similarity searches using the UniProtKB/Swiss-Prot database. Draft network assembly relied on protein-reaction associations available in the KEGG (Kyoto Encyclopedia of Genes and Genomes) database [28], with all reactions categorized as spontaneous or non-enzymatic also incorporated in the initial draft model. Hit selection was performed as described elsewhere [23] and phylogenetic proximity was implemented based on a phylogenetic tree from literature [22], this process automated via the “Automatic workflow” *merlin* tool and then integrated into the draft model [22].

### 2.3. Reversibility, Directionality and Balancing

Reaction reversibility and stoichiometry curation involved a multi-step process combining both automated and manual efforts. Initially, *merlin* was used to assist in correcting the direction and reversibility of reactions, utilizing references from remote databases like eQuilibrator [29] to predict reaction directionality, as described by Dias *et al.* [23]. This was followed by extensive manual curation, exploiting databases such as MetaCyc [30], Brenda [31], UniProt [32], FungiDB [33], RHEA [34], KEGG [30] and existing literature, in order to ensure that all reactions in the network are balanced, and with the correct directionality. All manually edited reactions can be found in Supplementary Data 1.

### 2.4. Compartmentalization and Transport reactions

This model includes four compartments: extracellular, cytoplasm, mitochondrion, and peroxisome and one intercompartment, the cytoplasmic membrane. The prediction of compartments for each enzyme was performed using the DeepLoc - 2.0 [35] and directly imported to *merlin*. The transport reactions were automatically generated by TranSyT [36], a tool integrated in *merlin*, based on the public database TCDB [37]. Additional transport reactions across internal and external membranes for common metabolites, such as H_2_O, CO_2_, and NH_3_, often carried out without a transporter, were added to the model with no gene association.

### 2.5. Biomass Equation

The biomass formation was depicted through an equation including proteins, DNA, RNA, lipids, carbohydrates, and cofactors, with detailed composition information for each macromolecule sourced from literature or experimental data. All calculations were performed as in previously described methodology [38] and are detailed in Supplementary Data 1. ATP requirements for biomass production and growth-associated maintenance (GAM) were added to the biomass equation with a value of 25.65 mmol ATP/gDCW, based on the ATP requirements for the biosynthesis of cell polymers as reported in [39], and ATP requirements for non-growth-associated maintenance (NGAM) was inserted in the model by an equation with specific fixed flux boundaries inferred from *Candida tropicalis* [39]. The theoretical phosphorus-to-oxygen ratio used in the *Saccharomyces cerevisiae* iMM904 metabolic model was applied to our model adding three generic reactions contributing to this ratio:

**Reaction R00081:**

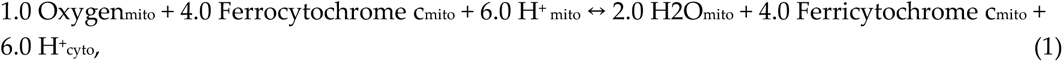

**Reaction R_Ubiquinol_Cytochrome_Reductase:**

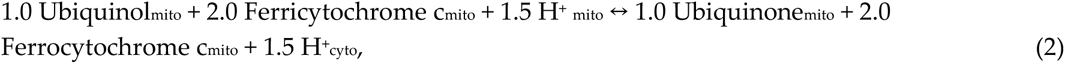

**Reaction T_ATP_Synthase:**

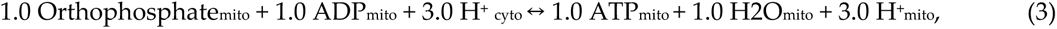

**The final balance reaction:**

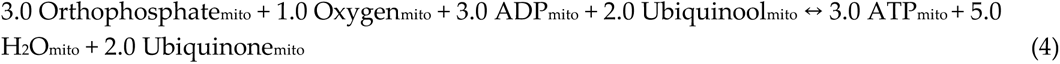

### 2.7 Network simulation and model curation

During the model reconstruction process, an extensive manual curation was needed in order to correct gaps in some pathways, due to incorrect reversibility, incomplete reactions, annotation errors, and blocked metabolites. Each case was meticulously inspected and studied, and reactions were edited, manually added to, or removed from the model based on evidence from the literature or deposited on databases such as KEGG pathways, MetaCyc, FungiDB etc. The detailed list of all the performed alterations can be found in Supplementary Data 1.

During this process, *merlin*’s “*Find blocked reactions*” was used to assist and accelerate the process. Additionally, *BioISO*, a tool based on the Constraints-Based Reconstruction and Analysis (COBRA) and Flux Balance Analysis (FBA) [40] frameworks, also integrated in *merlin*, assisted in the process of identifying potential errors in the network and accelerated the process of correcting the gaps.

### 2.8 Model Validation

#### 2.8.1. Strains and Growth Media

*C. neoformans var. grubii* H99E strain, from the laboratory of Jennifer Lodge was obtained from the Fungal Genomic Stock Center, and routinely maintained in Yeast extract–Peptone–Dextrose (YPD), containing: 20 g/L glucose (Merck, Darmstadt, Germany), 20 g/L peptone (Merck, Darmstadt, Germany), and 10 g/L yeast extract (Merck, Darmstadt, Germany). The parental KN99 and derived KN99_ΔCNAG_02553 were obtained from the deletion library created by the Madhani laboratory [41], through Fungal Genetics Stock Center, and grown on YNB medium, containing 1.7 g/L Yeast Nitrogen Base, without amino acids (Difco BD, England, United Kingdom) and 20 g/L inositol, used as carbon source. Synthetic minimal media (SMM) was used for batch cultivation experiments used to validate model predictions, SMM including: 20g/L glucose (Merck, Darmstadt, Germany), 2.7 g/L ammonium sulphate (Merck, Darmstadt, Germany), 0.05 g/L magnesium sulphate (Riddle-de-Haen), 2 g/L potassium dihydrogen phosphate (Panreac, Barcelona, Spain), 0.5 g/L calcium chloride (Merck, Darmstadt, Germany), and 100 µg/L biotin (Sigma).

#### 2.8.2. Aerobic Batch Cultivation

*C. neoformans var. grubii* cells were batch cultivated in Erlenmeyer flasks containing 250ml of SMM or YNB medium, at 30 °C (250 rpm). Exponential phase inocula, with an Optical Density (OD) (Hitachi u2001) at 600nm of 0.3, were prepared and cells were transferred to Erlenmeyer flasks containing 250ml of fresh medium and cultivated at 30 °C with orbital agitation (250 rpm) for the duration of the experiment.

#### 2.8.3. Cell Density, Dry Weight, and Metabolite Concentration Assessment

Throughout cell cultivation in SMM, 4 mL samples were collected every two hours for subsequent quantification of biomass and extracellular metabolites. Cell density was monitored by measuring OD_600nm_. For dry weight determination, culture samples were centrifuged at 13,000 rpm for 3 minutes, and the resulting pellets were freeze-dried for 72 hours at -80 ◦C before being weighed. Extracellular metabolites, including glucose, ethanol, glycerol, and acetic acid, were identified and quantified by High-performance liquid chromatography (HPLC) on an Aminex HPX-87 H Ion Exchange chromatography column, eluted with 0.0005 M H_2_SO_4_ at a flow rate of 0.6 mL/min at room temperature. Samples were analyzed in triplicate, and concentrations were determined using appropriate calibration curves. During the exponential growth phase, the specific growth rate, specific glucose consumption rate, and specific production rates of ethanol, glycerol, and acetic acid were calculated as described elsewhere [42].

#### 2.8.4. Network simulation and analysis

All the phenotype simulations were performed with Flux Balance Analysis (FBA) in OptFlux 3.0 using the IBM”CPLEX solver, including: gene and reaction essentiality; growth assessment; metabolite production and consumption; and carbon and nitrogen source utilization. For gene and reaction essentiality, *in silico* growth was simulated in environmental conditions mimicking RPMI medium and a biomass flux lower than 5% of the wild-type strain, after the respective gene/reaction knockout, was considered the threshold for essentiality. Gene and reaction knockout was simulated by restraining its corresponding flux bounds to zero.

## 3. Results and Discussion

### 3.1. Model characteristics

The *C. neoformans var. grubii* genome-scale metabolic model reconstructed herein, and denominated iRV890, comprises 890 genes associated with 2598 reactions, of which 683 are transport reactions, and 2047 metabolites across four compartments (extracellular, cytoplasm, mitochondria, and peroxisome). The model can be found in SBML format in Supplementary Data 2. Among the 2598 reactions, 1747 are cytoplasmic, 351 mitochondrial, 60 peroxisomal and 440 are drains (exchange constraints, used to simulate the import of media components or the leakage or export of extracellular metabolites).

During the manual curation process, a total of 639 reactions/genes required alterations, including 80 who were mass balanced, 518 who were corrected for reversibility, directionality, or added or removed from the model, and 41 whose annotation was corrected, as detailed in Supplementary Data 1.

The Biomass equation (Table 1) encompasses the cell’s major components along with their respective and relative contributions, including DNA, RNA, lipids, carbohydrates, and cofactors. The equation’s composition in carbohydrates [43], and lipids [44–46] was inferred from literature data for *C. neoformans*. The composition of Proteins, DNA and RNA was determined by the e-BiomassX where the whole genome sequence was used to estimate the amount of each deoxyribonucleotide as described in [47] mRNA, rRNA, and tRNA being used to estimate the total RNA in the cell as described in [47,48].

**Table 1:** Biomass composition used in the model iRV890. The full individual validated contributions of each of these metabolites are shown in Supplementary Data 1.

| Metabolite | g/gDCW | Metabolite | g/gDCW |
| --- | --- | --- | --- |
| **Lipids** |  | **Proteins** |  |
| Lanosterol | 0.000122 | L-Valine | 0.019058 |
| Zymosterol | 0.000254 | L-Tyrosine | 0.020501 |
| Squalene | 0.000209 | L-Tryptophan | 0.006392 |
| Ergosterol | 0.000724 | L-Threonine | 0.022013 |
| Phosphatidylserine | 0.005024 | L-Serine | 0.027696 |
| Phosphatidylinositol | 0.004638 | L-Proline | 0.015390 |
| Phosphatidylcholine | 0.031241 | L-Phenylalanine | 0.022928 |
| Phosphatidylethanolamine | 0.017714 | L-Methionine | 0.008274 |
| Cardiolipin | 0.002254 | L-Lysine | 0.033668 |
| Phosphatidic acid | 0.000644 | L-Leucine | 0.036895 |
| Phosphatidylglycerol | 0.000644 | L-Isoleucine | 0.028492 |
| Tetradecanoic acid | 0.000020 | L-Histidine | 0.010159 |
| Hexadecanoic acid | 0.000097 | L-Glutamate | 0.029371 |
| Octadecanoic acid | 0.000038 | L-Cysteine | 0.003902 |
| Dodecanoic acid | 0.000021 | L-Aspartate | 0.023883 |
| Decanoic acid | 0.000011 | L-Asparagine | 0.027060 |
| Octanoic acid | 0.000038 | L-Arginine | 0.020979 |
| Octadecanoic acid | 0.000038 | L-Alanine | 0.012706 |
| (9Z)-Octadecenoic acid | 0.000093 | Glycine | 0.010258 |
| (9Z,12Z)-Octadecadienoic acid | 0.000116 | L-Glutamine | 0.020550 |
| (9Z,12Z,15Z)-Octadecatrienoic acid | 0.000002 |  |  |
| Triacylglycerol | 0.032969 | **Soluble Pool** |  |
| Sterol esters | 0.001127 | Pyridoxine 5'-phosphate | 0.000833 |
|  |  | FAD | 0.000833 |
| **Carbohydrates** |  | Thiamine(1+) diphosphate | 0.000833 |
| Chitin | 0.005645 | NAD | 0.000833 |
| Mannan | 0.033956 | Glutathione | 0.000833 |
| $\beta$ (1,3)-Glucan | 0.360399 | Riboflavin | 0.000833 |
|  |  | Eumelanin | 0.000833 |
| **Ribonucleotides** |  | Ubiquinone-6 | 0.000833 |
| UTP | 0.006713 | NADP | 0.000833 |
| GTP | 0.006806 | COA | 0.000833 |
| CTP | 0.005381 | FMN | 0.000833 |
| ATP | 0.007101 | 5-Methyltetrahydrofolate | 0.000833 |
| **Deoxyribonucleotides** |  |  |  |
| dTTP | 0.016718 |  |  |
| dGTP | 0.017029 |  |  |
| dCTP | 0.015059 |  |  |
| dATP | 0.017193 |  |  |

The translated genome sequence was used to calculate the amino acid composition using the percentage of each codon usage as described in [47]. Essential metabolites were included in the biomass composition to qualitatively account for the essentiality of their synthesis pathways [49,50]. The growth and non-growth ATP requirements were adopted from *S. cerevisiae* [51].

### 3.2. Model validation

#### 3.2.1 Carbon and nitrogen source utilization

*In silico* simulations were conducted using 222 different compounds as the exclusive carbon or nitrogen sources, under conditions mimicking those of the minimal medium reported in [21]. The *in silico* growth was compared to publicly available phenotypic microarray (Biolog platform) data for *C. neoformans var. grubii* performed in [21]. A total of 155 sole carbon sources and 67 sole nitrogen sources were evaluated. For the analyses we used the data from stationary phase yeasts condition after calculating the difference from the respective negative control group, without any carbon or nitrogen sources. iRV890 model correctly predicted growth in 85% (133/155) of the carbon sources tested and in 85% (57/67) of the nitrogen sources Supplementary Table 1. In some cases of failed predictions, such as L-ornithine and glycerol (carbon source) and amino acids and D-Glucosamine (nitrogen source), genetic information and the model include all the necessary steps to predict their assimilation as sole carbon/nitrogen sources, but no growth was experimentally observed. In such cases, the failed prediction may be related to non-metabolic factors that are not considered in model simulations, or to inaccuracies regarding the annotation of transporters, which is still a big challenge in the current model development process [52]. In other cases, however, the prediction model failed because specific enzymes are not yet characterized for *C. neoformans*, despite growth in experimental conditions. The comparison between the model’s prediction and experimental evidence suggests that the following enzyme activities are likely to be present in *C. neoformans*, although the underlying genes and proteins were not yet identified: 1.2.1.3 (aldehyde dehydrogenase), 1.1.1.21 (aldose reductase), 3.1.1.65 (L-rhamnono-1,4-lactonase), 1.1.1.56 (ribitol 2-dehydrogenase), 5.1.3.30 (D-psicose 3-epimerase), 2.7.1.55 (allose kinase), 4.1.2.10 ((R)-mandelonitrile lyase), 5.3.1.3 (D-arabinose isomerase), 3.2.1.86 (6-phospho-beta-glucosidase), 4.1.2.4 (deoxyribose-phosphate aldolase), 3.2.1.86 (6-phospho-beta-glucosidase) and 1.1.1.16 (galactitol 2-dehydrogenase). The identification and characterization of these predicted functions and their underlying gene(s) will shed light on the specific pathways of carbon or nitrogen assimilation in this pathogen, potentially revealing new mechanisms of virulence related to adaptation to the host environment. Altogether, the model achieved 85% predictability which is a high value, especially considering that the extensive list of carbon and nitrogen sources tested includes many that are not commonly used in traditional metabolic and phenotypic experiments and thus lack biochemical characterization.

#### 3.2.2 Growth parameters in batch culture

To quantitatively validate the model, the specific growth rate, glucose consumption rate, and metabolite production rates were experimentally determined, and compared with *in silico* predicted values. For a glucose consumption rate of 1.72 mmol.gDCW^-1^.h^-1^, a specific growth rate of 0. 188 h^-1^ was experimentally determined, leading to no detectable production of ethanol, glycerol, or acetate. For comparison with *in silico* results, we simulated the system’s behavior in SMM medium with a fixed glucose uptake flux of 1.72 mmol.g^-1^ dry weight.h^-1^. Other nutrient fluxes were left unconstrained, as the system was glucose-limited under these conditions. The simulation predicted a specific growth rate of 0.128 h^-1^, a difference of 0.06 h^-1^ to the experimentally determined value (Table 2). In these conditions, the model did not predict the formation of glycerol, acetic acid, or ethanol as by-products, consistent with the experimental data. Moreover, the model is accurate at predicting no growth of *C. neoformans* under anaerobic conditions which is to be expected since this pathogen is an obligate aerobic fungus [53].

**Table 2.** Growth parameter values predicted by the iRV890 model, in comparison with those determined experimentally.

|  | Specific growth rate (h<sup>-1</sup>) | q (mmol g<sup>-1</sup> dry weight h<sup>-1</sup>) |  |  |  |
| --- | --- | --- | --- | --- | --- |
|  |  | Glucose | Ethanol | Glycerol | Acetic acid |
| *In silico* | 0.128 | 1.72 | 0 | 0 | 0 |
| *In vivo* | 0.188 | 1.72 | 0 | 0 | 0 |

### 3.3. *C. neoformans* unique metabolic features

To uncover unique metabolic features of this pathogen, a comparison was made between the *C. neoformans* GSMM with those previously built for *C. glabrata* [49], *C. albicans* [54], *C. auris* [55] and *S. cerevisiae* [56] by us and others. A comparison across the existing models was carried out based on shared EC numbers. After intersecting the EC numbers present in each of the five models, 40% (229/566) of the EC numbers were common among all the tested yeasts (Figure 1). Additionally, the remaining 17% (96/556) are exclusive to the *C. neoformans* model and may represent unique metabolic features of this species relative to the remaining. We confirmed none of these 96 EC numbers were associated with outdated, incomplete or incorrect reaction associations. However, a small subset of these 96 EC numbers may be present in other species included in the comparison, but not accounted for in their respective GSMMs during the process of reconstruction.

**Figure 1:**
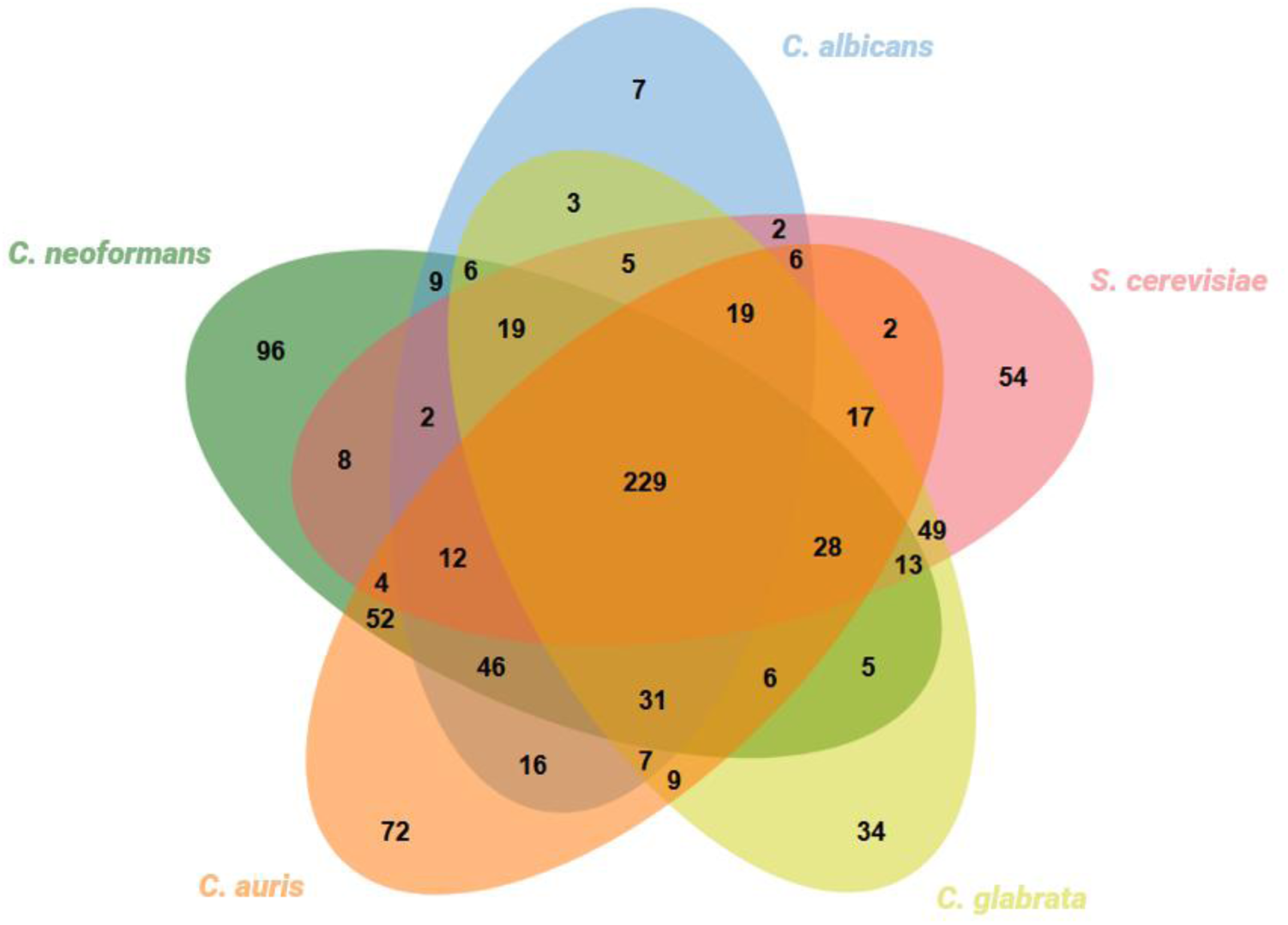
Multi-species comparison in terms of proteins with an associated EC number present in the *C. neoformans* iRV890, *C. albicans* iRV781, *C. auris* iRV973, *S. cerevisiae* iIN800 and *C. glabrata* iNX804 GSMMs. The multiple intersection was performed using jvenn [57].

From the list of 96 *C. neoformans* unique EC numbers (Supplementary Data 1), metabolic features or pathways relevant in the context of fungal infection in the host brain were searched manually for, and compared to extant knowledge of these pathways being defense mechanisms, or enabling host adaptation, through degradation or biosynthesis of specific metabolites. A few of these unique EC numbers with higher potential of impacting *C. neoformans* pathogenesis are discussed below:

**1.1.1.12 and 1.1.1.287-** L-arabinitol 4-dehydrogenase and D-arabinitol dehydrogenase are two enzymes that are required for L-arabinitol assimilation as carbon source, which is a particular metabolic feature of *C. neoformans* when compared to other yeast species (Supplementary Table 1). Indeed, neither *Candida* species [54,58] nor *S. cerevisiae* [59], can assimilate L-arabitol, unless genetically engineered [60]. Interestingly, environment isolates containing SNPs in the *PTP1* gene, encoding a *C. neoformans* arabitol transporter, were associated with increased patient survival, while a virulence defect was observed in BALB/c mice due to *PTP1* gene deletion [61]. *PTP1* expression was also found to be highly induced in macrophage and amoeba infection [62]. Since arabitol is present in the cerebrospinal fluid [63], it is possible this pathway may be used to feed from polyols in CNS and contributes to explain brain tropism of *C. neoformans*, compared to other fungal species.

**1.1.3.8 and 3.1.1.17 -** L-gulonolactone oxidase and gluconolactonase are two enzymes that participate in ascorbate metabolism, allowing the utilization of Inositol and D-glucuronate as source for L-ascorbate biosynthesis (Figure 2). Interestingly, it was reported by two independent studies that the presence of ascorbate, an antioxidant, lowers the susceptibility towards fluconazole in *C. neoformans* [64,65]. However, this effect seems to not be related to its antioxidant potential, but with ascorbate-induced up-regulation of Upc2, a transcriptional regulator of genes involved in ergosterol biosynthesis, as shown in *C. albicans* [66]. The ability of *C. neoformans* to synthesize ascorbate from inositol is particularly noteworthy, given the abundance of inositol in the human brain [20] and the widespread use of fluconazole in treating infections. Further it is possible that ascorbate contributes to resistance to ROS. Having a mechanism to produce a compound that mitigates the toxicity of fluconazole and ROS could contribute to a significant adaptive advantage for this species.

**Figure 2.**
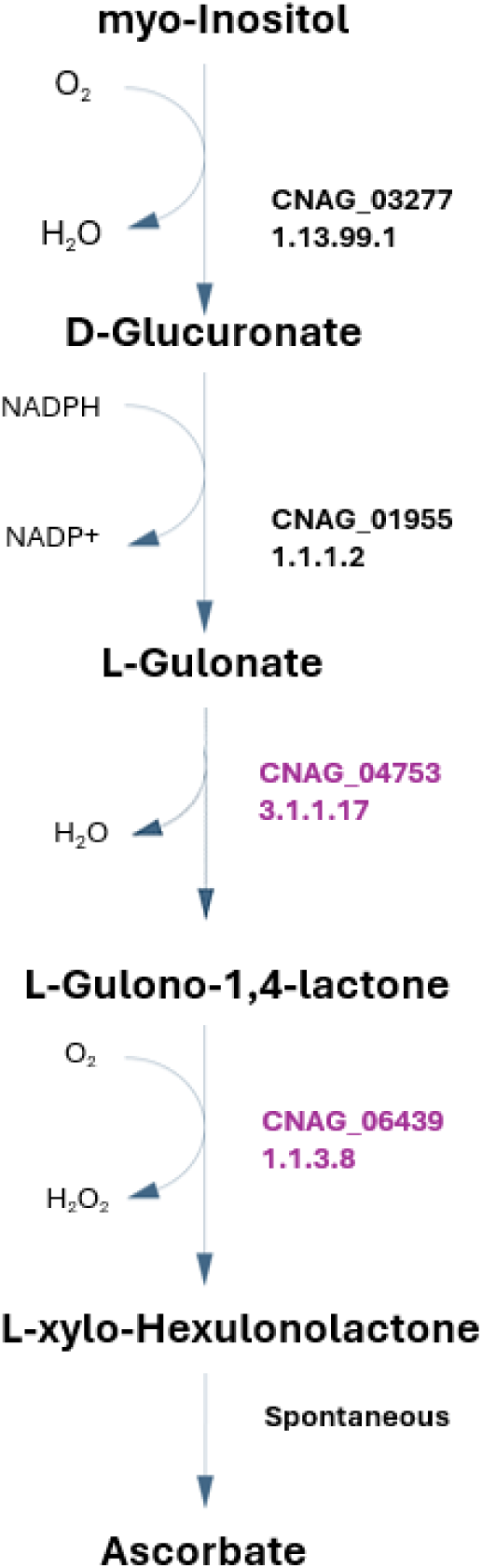
*C. neoformans* pathway for ascorbate biosynthesis, with the respective *C. neoformans var. grubii* EC numbers present in the iRV890 model. The 1.1.3.8 and 3.1.1.17 enzymes, which are unique to *C. neoformans* among other pathogenic yeasts, are highlighted in purple.

Additionally, the 1.1.3.8 and 3.1.1.17 enzymes are also important for inositol assimilation as a carbon source through a variation of the previous pathway. This pathway was suggested recently as an alternative pathway in fungi for inositol assimilation, and since inositol is highly abundant in the human brain, this may represent a very important metabolic feature for *C. neoformans*. In fact, in order to implement that pathway in the model, two of the reactions reported were recreated and attributed with the names R2_Inositol_Pathway and R1_Inositol_Pathway in the model, although the corresponding EC numbers and genes have not been identified in the annotated *C. neoformans* genome [67]. This pathway was recreated exclusively from literature, and while it lacks validation studies, two possible genes were hypothesized as probable candidates for encoding the 1.1.1.69 enzyme, CNAG_02553 and CNAG_00126, predicted by OrthoMCL [68]. Additional pathways for inositol assimilation are reported for animal (Figure 3.B) and bacteria (Figure 3.C); however, since *C. neoformans* lacks almost all the enzymes in those pathways, we considered that the new pathway reported in fungi [67] was the most probable to occur in this pathogen. Taking advantage of the available ΔCNAG_02553 deletion strain, we tested whether a strain deleted for this putative enzyme could be grown in inositol as a single carbon source, compared to parental strain. However, even in the absence of CNAG_02553 gene, *C. neoformans* is able to utilize inositol as the sole carbon source in SMM (YNB, supplemented with glucose or inositol, data not shown). Eventually, it would be necessary to knockout both CNAG_00126 and CNAG_02553 genes to obtain a strain unable to grow in media containing inositol as the sole carbon source. Further scrutiny is required to address this issue.

**Figure 3.**
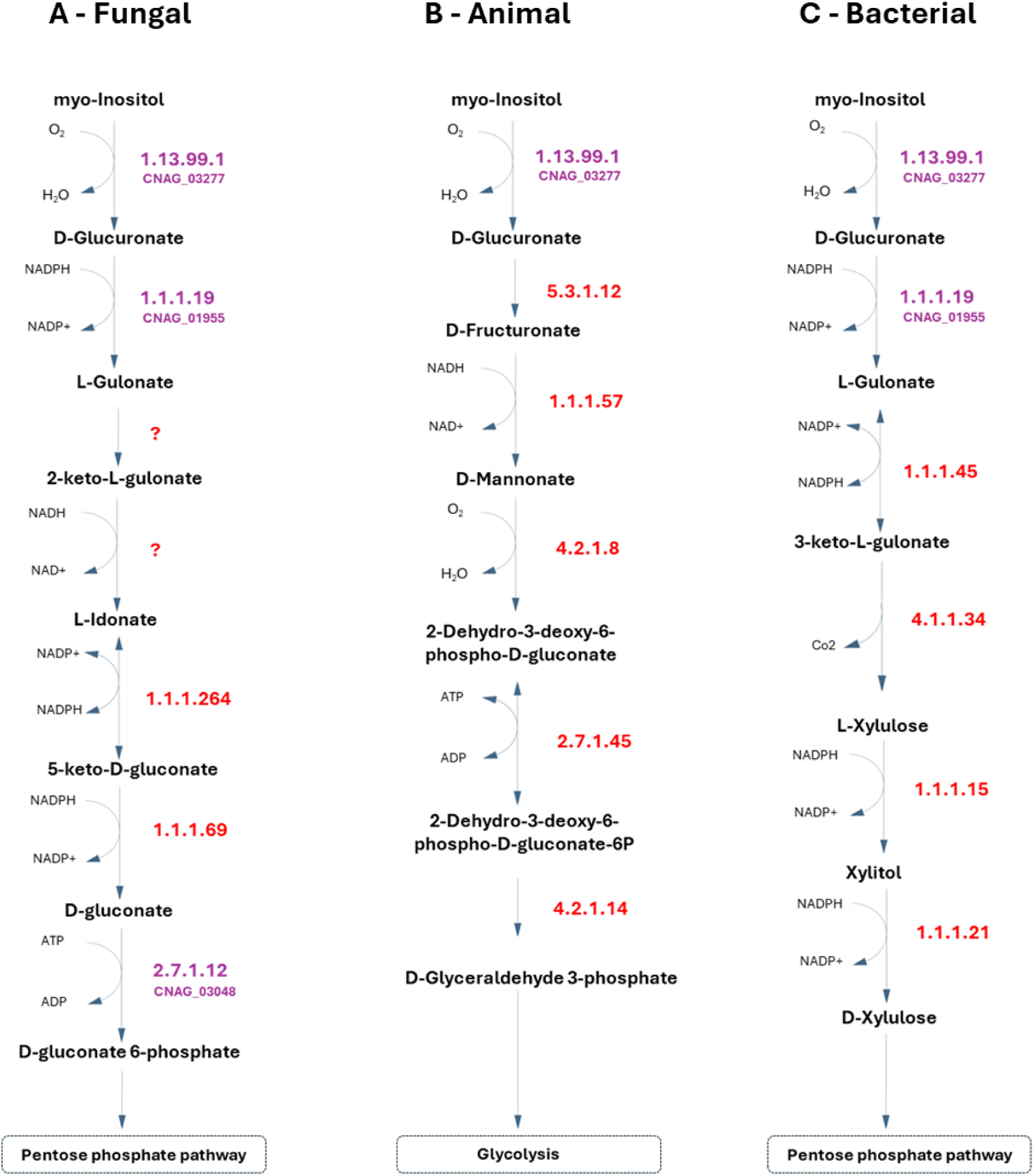
Metabolic pathways for inositol assimilation as carbon source, A - based on the proposed fungal inositol assimilation pathway reported in Kuivanen et al. 2016 [67], B – based on the animal inositol assimilation pathway, C-based on the bacterial inositol assimilation pathway, and D- the unknown, and apparently unique, pathway of inositol assimilation proposed for *C. neoformans*. The respective *C. neoformans var. grubii* genes present in iRV890 are highlighted in purple. The currently unknown genes are highlighted in red and the proposed reactions with an unknown EC number are represented as a question mark in red.

**1.1.1.377 -** L-rhamnose 1-dehydrogenase is required for L-rhamnose assimilation as sole carbon source. Rhamnose is used by some pathogens, for example *Pseudomonas aeruginosa*, to produce rhamnolipids, and constitutes an important virulence factor in those bacteria, with roles in biofilm formation, hydrophobic nutrient uptake, and host immunity evasion, characterized for increasing lung epithelial permeability [69,70] and inhibition of macrophage phagocytosis [71]. *Candida* species [54,55,58] and *S. cerevisiae* (unless engineered) [72] cannot assimilate L-rhamnose, and thus assimilation of rhamnose is a particular metabolic feature of *C. neoformans* when compared to these yeast species (Supplementary Table 1).

**1.14.13.231 -** tetracycline 11a-monooxygenase is an enzyme that allows the direct conversion of tetracycline into 11a-hydroxytetracycline, reported to confer resistance to all clinically relevant tetracyclines, by efficient degradation of a broad range of tetracycline analogues. The hydroxylated product, 11a-hydroxytetracycline, is very unstable and leads to intramolecular cyclization and non-enzymic breakdown to undefined products, completely neutralizing the tetracycline effects [73,74]. Although tetracyclines are generally used as antibacterial antibiotics, and have poor antifungal activity, the presence of this enzyme in *C. neoformans* should be taken into consideration when designing tetracyclines against fungi.

**3.1.3.8 -** 3-phytase is an enzyme involved in inositol metabolism that may be involved in the production of phytic acid from inositol, a primary storage molecule of phosphorus and inositol in fungi (although not in the pathogenic *Candida* species), bacteria and plants [75]. Interestingly, this pathway has been shown to play a key role in *C. neoformans* virulence. Indeed, it was previously reported that the deletion of the gene encoding the enzyme (EC number 2.7.1.158) that immediately precedes 3-phytase leads to growth impairment and to attenuated virulence in *C. neoformans*, associated with failed dissemination into the brain [76].

**3.5.2.17 –** hydroxyisourate hydrolase is an enzyme essential for the assimilation of uric acid as sole nitrogen source. Uric acid is a normal component of urine and bird guano. In bird guano, 70% of the nitrogen present is in the form of uric acid with the rest consisting primarily of xanthine, urea, and creatinine [77] Additionally, uric acid enhances the production of key cryptococcal virulence factors, including capsule and urease, an enzyme required for full fitness at mammalian pH and dissemination to the brain [78], *C. neoformans* capsule is induced in the presence of uric acid [79].

**4.1.1.105 –** L-tryptophan decarboxylase catalyzes the conversion of L-tryptophan into Tryptamine, which can then be converted into serotonin, and shares structure with several aminergic neuromodulators. However, the reaction is bidirectional, and Tryptamine can also be converted into L-tryptophan. While it is unclear which may be the role of this enzyme in *C. neoformans*, it is potentially related to the brain environment, specifically in the utilization of serotonin as a nitrogen source, through its conversion to L-tryptophan.

**4.1.1.28, 1.14.18.1, and 1.10.3.2 –** DOPA decarboxylase, tyrosinase and laccase are particularly important in *C. neoformans*, as they are involved in the biosynthesis of melanin. Most fungi possess multiple melanin biosynthetic pathways, while *Cryptococcus neoformans* exclusively synthesizes melanin through the L-DOPA pathway. [80]. Melanin is able to neutralize oxidative stress radicals as well as protecting the pathogen against the host immune system and antifungal drugs, such as caspofungin and amphotericin B. L-DOPA and Dopamine are present in the human brain and serve as precursors for dopamine biosynthesis in this pathogen, but it is uncertain why *C. neoformans* exclusively uses this pathway, compared to other human pathogenic fungi.

### 3.4. Drug target analysis based on gene essentiality prediction

Pathogen’s GSMM are particularly useful to identify potential new drug targets, among predicted essential genes. For that purpose, a list of all predicted essential genes and enzymes in *C. neoformans* was obtained through simulation of the system’s behaviour in RPMI medium, which mimics the environmental conditions of human serum. A total of 157 enzymes and 101 genes were identified as essential in RPMI medium. Among these targets, some have been previously identified as essential genes in other pathogenic yeasts (see Table 3), indicating potential drug targets common to all *Candida* species and *C. neoformans*. Notably, Erg11 and Fks1 are already targets of currently used antifungals, fluconazole and echinocandins, respectively. Additionally, Erg26, Erg27, Erg24, Erg4, Erg7, Erg12, and Erg13 have all been identified herein as potential drug targets within the ergosterol biosynthetic pathway. Particularly interesting is Erg4, as it lacks a human ortholog, and may represent a superior candidate for designing compounds with enhanced selectivity and lower toxicity.

**Table 3.**
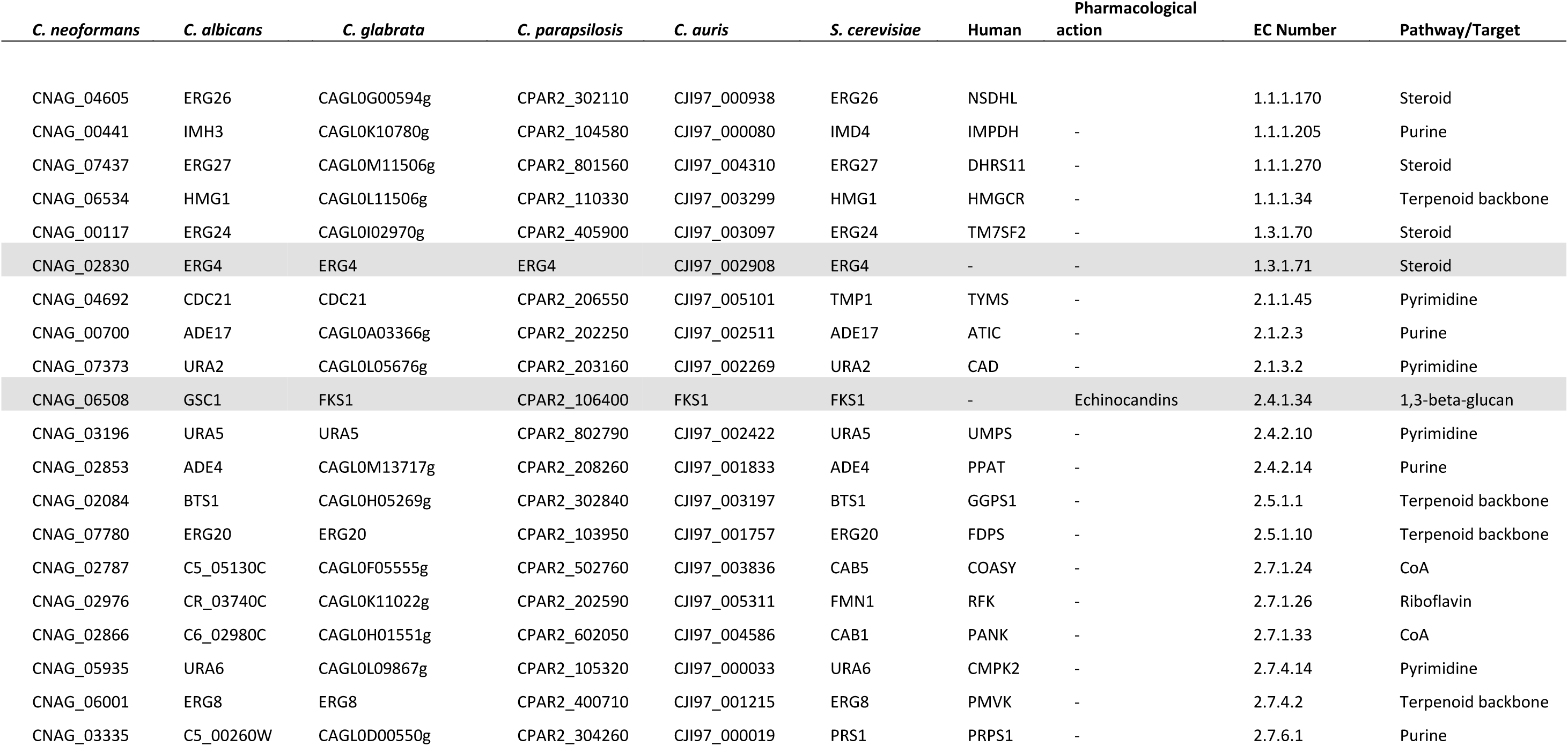
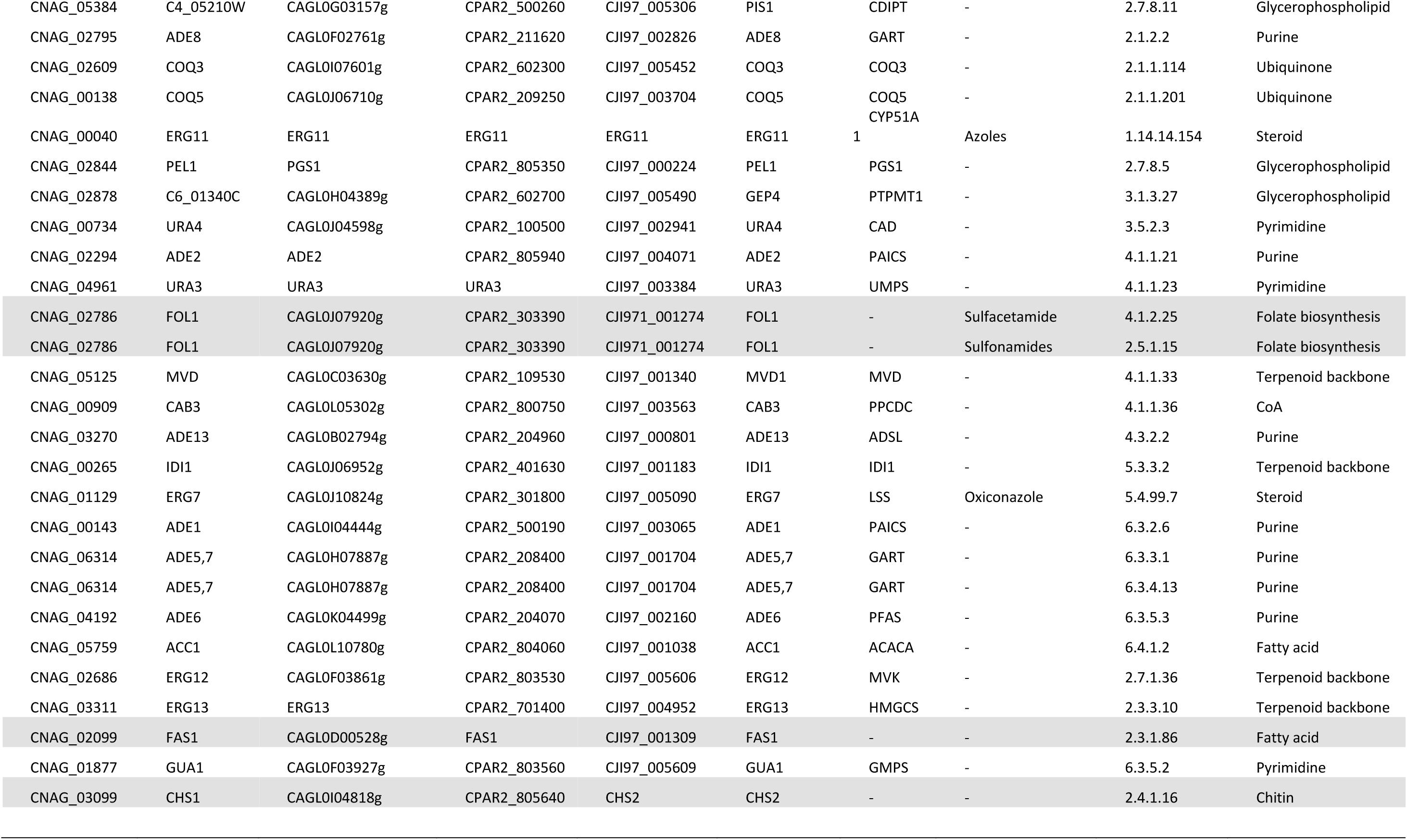
Enzymes predicted to be essential in RPMI medium in 5 pathogenic fungal species, based on the screening of the genome-scale metabolic models of *C. neoformans* iRV890, *C. auris* IRV973, *C. parapsilosis* iDC1003, *C. albicans* iRV781, and *C. glabrata* iNX804. Grey rows highlight enzymes which are not encoded in the human genome. Data regarding the drug association retrieved from the DrugBank database; only drugs with known pharmacological action against pathogens were selected.

Similarly to *Candida*, which lacks a Folate transporter [81] and relies on its *de novo* biosynthesis, *C. neoformans* seems to also lack a folate transporter, leading to the identification of Fol1, a multifunctional enzyme of the folic acid biosynthesis pathway, as a promising multi-yeast drug target. Furthermore, Fas1, a fatty acid synthase enzyme, and Chs1, a chitin synthase, also lack human orthologs and constitute promising alternative antifungal drug targets due to their important role for membrane and cell wall structure and integrity. Other noteworthy targets span various pathways, including purine metabolism, terpenoid backbone biosynthesis, pyrimidine metabolism, CoA biosynthesis, glycerophospholipid biosynthesis, and ubiquinone biosynthesis (Table 3). However, exploring these targets requires leveraging potential structural differences in the enzyme’s active site compared to their human counterparts.

Since *C. neoformans* colonizes a different host environment and is phylogenetically distant from *Candida* spp. our evaluation was extended to include potential new drug targets that are unique to this species, and not shared by *Candida* spp. We identified only two such targets: the 1.14.18.1 tyrosinase, encoded by the gene CNAG_03009, and the 2.5.1.83 hexaprenyl diphosphate synthase, encoded by the gene CNAG_04375. While tyrosinase, responsible for melanin production, has a human ortholog (since humans also synthesize melanin via the L-DOPA), hexaprenyl diphosphate synthase (2.5.1.83) is fungal-specific and may represent an interesting target. This enzyme plays a crucial role in terpenoid backbone biosynthesis, serving as a key contributor to the synthesis of precursors for ubiquinone biosynthesis.

## 4. Conclusions

The construction and validation of iRV890, the first genome-scale metabolic model for *C. neoformans var. grubii* is presented herein. iRV890 constitutes a robust platform for exploring and elucidating the metabolic features of this poorly understood pathogen, particularly concerning its interaction within the central nervous system and the human host. By encompassing 890 genes associated with 1466 reactions, this model offers a comprehensive view of the metabolic landscape of the pathogen. Through *in silico* simulations, we predicted the use of more than 200 compounds as sole carbon or nitrogen sources, and after comparison to experimental data from phenotypic microarrays we gained valuable insights into the metabolic capabilities of *C. neoformans*. The model correctly predicts 85% of the sole carbon and nitrogen sources tested. The model was able to accurately predict the organism’s specific growth rate and confirmed its inability to grow under anaerobic conditions or to accumulate glycerol, acetic acid, or ethanol as metabolic by-products during growth in synthetic minimal medium, with glucose as carbon source. Additionally, we propose a list of yet unidentified enzymes expected to be present in *C. neoformans*, based on the carbon and nitrogen utilization and with potential to represent new host adaptation or virulence mechanisms, including new clues on the pathway for inositol utilization in *C. neoformans*.

Our investigation into the unique metabolic features of *C. neoformans* has unveiled several pathways and enzymatic activities that are proposed to play pivotal roles in fungal infection within the host brain. Some enzymes constitute important virulence factors, such as Tyrosinase and laccase, enzymes responsible for production of melanin which has an important role in host immune evasion [15], infection proliferation and drug resistance [16,17]. Other enzymes are related to drug and stress resistance, such as tetracycline 11a-monooxygenase, L-gulonolactone oxidase and gluconolactonase. The remaining enzymes are directed related to alternative carbon/nitrogen source utilization and are important for environmental adaptation.

For example, hydroxyisourate hydrolase is essential for the assimilation of uric acid as a nitrogen source, an important virulence factor mechanism, and 3-phytase is involved in inositol metabolism and storage, important for brain dissemination.

In this work we also propose several potential drug targets in *C. neoformans*. Notably, enzymes such as Erg4, Chs1, Fol1 and Fas1 present promising opportunities for targeted drug development, due to their absence in human cells, offering opportunities for the development of selective and low-toxicity compounds. The CNAG_03009 and CNAG_04375 genes, encoding a tyrosinase and a hexaprenyl diphosphate synthase, are presented as potential antifungal drug targets specific to C. *neoformans*.

Our model contributes to a better understanding of *C. neoformans* metabolism, especially within the host environment. With this work, we not only propose new metabolic enzymes awaiting characterization but also offer insights into key pathways and interactions shaping the dynamics between host and pathogen and its adaptive strategies. We also propose some potential antifungal targets for *C. neoformans* and confirmed the coverage of already identified targets also to that species. These results hold promise for the discovery of novel drug targets and for the full comprehension of this pathogen’s metabolic network with an expected impact in combating cryptococcosis.

## Author contributions

R.V., C.C., and M.C.T. conceived and designed the study. R.V. performed the model construction & development, data analysis and curation. R.V., D.C., and W.N. performed the experiments with *C. neoformans*. O.D. contributed to model construction and data analysis. L.C. performed data analysis. R.V. wrote the original draft preparation. R.V., L.C., C.C., and M.C.T. reviewed and edited the final version. All authors have read and agreed to the published version of the manuscript.

## CRediT authorship contribution statement

**Romeu Viana:** Writing – review & editing, Writing – original draft, Visualization, Methodology, Investigation, Formal analysis, Data curation. **William Newton**: Methodology, Investigation. **Diogo Couceiro:** Methodology, Investigation. **Oscar Dias**: Methodology, Formal analysis. **Luís Coutinho:** Writing – review & editing, Formal analysis. **Carolina Coelho:** Writing – review & editing, Supervision, Methodology, Conceptualization. **Miguel Cacho Teixeira:** Writing – review & editing, Supervision, Methodology, Conceptualization, Funding.

## Acknowledgements

The authors acknowledge the OSCARS project, funded by the European Commission’s Horizon Europe Research and Innovation Programme under grant agreement No. 101129751. This work was further financed by national funds from “Fundação para a Ciência e a Tecnologia” (FCT) (AEM PhD grant to RV; projects UIDB/04565/2020 and UIDP/04565/2020 of the Research Unit Institute for Bioengineering and Biosciences—iBB; project UIDB/04469/2020 for the Centre of Biological Engineering—CEB; project LA/P/0029/2020 for LABBELS –Associate Laboratory in Biotechnology, Bioengineering and Microelectromechanical Systems; and project LA/P/0140/2020 for the Associate Laboratory Institute for Health and Bioeconomy—i4HB). WN was funded by a DTP BRC Exeter NIHR203320. Fungal strain collection was funded via NIH funding (R01AI100272) led by Hiten Madhani, UCSF. This work was supported by AMS Springboard Award SBF006\1024 (UK), and Wellcome Trust Institutional Strategic Support Award (WT105618MA) to C.C. We acknowledge other funding from the MRC Centre for Medical Mycology at the University of Exeter (MR/N006364/2 and MR/V033417/1). This study/research is funded by the National Institute for Health and Care Research (NIHR) Exeter Biomedical Research Centre (BRC). The views expressed are those of the author(s) and not necessarily those of the NIHR or the Department of Health and Social Care.

## Appendix A. Supplementary material

