## Appendix A for "Reconstruction and exploitation of a dedicated Genome-Scale Metabolic Model of the human pathogen *C. neoformans*": Appendix_A.Suplementary_material.docx

### Appendix A. Supplementary material

**Supplementary Data 1 -** 1 and 2: Detailed information regarding biomass composition of *Cryptococcus neoformans* used in iRV890.	3, 4 and 6: Detailed information regarding model iRV890 curation process. 6: List of essential enzymes in RPMI medium environmental conditions in *C. neoformans* iRV890. 7: List of all EC numbers present in *C. neoformans* iRV890, *C. parapsilosis* iDC1003, *C. albicans* iRV781, *C. glabrata* iNX804 and *S. cerevisiae* iIN800, and respective C. neoformans iRV890 unique EC numbers, with detailed description.

https://drive.google.com/drive/folders/1oZUtNrh9nMAtFv7qcjiNYZrw3L6ZU5ty?usp=sharing

**Supplementary Data 2 -** Model iRV890 in sbml format.

https://drive.google.com/drive/folders/1oZUtNrh9nMAtFv7qcjiNYZrw3L6ZU5ty?usp=sharing

**Supplementary Table 1 -** Comparison between experimental and *in silico* phenotypic behavior of *C. neoformans* under different carbon and nitrogen sources. Highlighted in grey are the cases that are not in accordance with both evidence. Carbon source utilization was predicted correctly in 86% (133/155) of the cases and nitrogen source 85% (57/67). Growth (+); lack of growth (–);

| **Carbon source** | ***In silico*** | **Experimental** |
| --- | --- | --- |
| Acetamide | - | - |
| Acetic acid | + | + |
| Acetoacetate | - | - |
| Adenosine | - | - |
| a-D-Glucose | + | + |
| a-D-Lactose | - | - |
| a-Hydroxyglutaric acid-g-Lactone | - | - |
| a-Keto-Valeric acid | - | - |
| Amygdalin | - | + |
| Arbutin | - | + |
| b-Hydroxybutyric acid | - | - |
| b-Methyl-D-Galactoside | - | - |
| b-Methyl-D-Xyloside | - | - |
| Bromosuccinic acid | - | - |
| Butylamine | - | - |
| Butyric acid | - | - |
| Capric acid | - | - |
| Caproic acid | - | - |
| Citric acid | - | - |
| D,L-Carnitine | - | - |
| D,L-Malic acid | - | - |
| D-Alanine | - | - |
| D-Allose | - | + |
| D-Arabinose | - | + |
| D-Arabitol | + | + |
| D-Aspartic acid | - | - |
| D-Cellobiose | + | + |
| Deoxyadenosine | - | - |
| Deoxyribose | - | + |
| Dextrin | + | + |
| D-Fructose | + | + |
| D-Fructose-6-Phosphate | - | - |
| D-Galactarate | - | + |
| D-Galactose | + | + |
| D-Galacturonate | - | + |
| D-Glucarate | - | - |
| D-Gluconic acid | + | + |
| D-Glucosamine | + | + |
| D-Glucose-1-Phosphate | - | - |
| D-Glucose-6-Phosphate | - | - |
| D-Glucuronate | + | + |
| D-Lactic acid Methyl Ester | - | - |
| D-Lactitol | - | - |
| D-Malic acid | - | - |
| D-Mannitol | + | + |
| D-Mannose | + | + |
| D-Melibiose | - | - |
| D-Psicose | - | + |
| D-Raffinose | + | + |
| D-Ribose | + | + |
| D-Serine | + | - |
| D-Sorbitol | + | + |
| D-Tagatose | - | + |
| D-Tartaric acid | - | - |
| D-Threonine | - | - |
| D-Trehalose | + | + |
| D-Xylose | + | + |
| Ethanolamine | - | - |
| Formate | - | - |
| Fumaric acid | - | - |
| Galactitol | - | + |
| Gelatin | - | - |
| g-Hydroxybutyric acid | - | - |
| Glucuronamide | - | - |
| Glycerol | + | - |
| Glycerone phosphate | + | + |
| Glycine | - | - |
| Glycolate | - | - |
| Glyoxylate | - | - |
| Hydroxyproline | - | - |
| Inosine | - | - |
| Inulin | - | - |
| Itaconic acid | - | - |
| Lactulose | - | - |
| L-Alanine | + | + |
| L-Arabinose | + | + |
| L-Arabitol | + | + |
| L-Arginine | - | - |
| L-Asparagine | - | - |
| L-Aspartate | + | - |
| L-Fucose | - | - |
| L-glutamate | + | + |
| L-Glutamine | + | - |
| L-Gulono-1,4-lactone | + | + |
| L-Histidine | - | - |
| L-Homoserine | + | - |
| L-Isoleucine | - | - |
| L-Leucine | - | - |
| L-Lysine | - | - |
| L-Lyxose | + | + |
| L-Malic acid | - | - |
| L-Methionine | - | - |
| L-Ornithine | + | - |
| L-Phenylalanine | - | - |
| L-Proline | + | + |
| L-Rhamnose | - | + |
| L-Serine | + | - |
| L-Sorbose | + | + |
| L-Threonine | + | - |
| L-Valine | - | - |
| Malonate | - | - |
| Maltose | + | + |
| Mannan | - | - |
| Methyl beta-D-galactoside | - | - |
| Methyl ethyl ketone | - | - |
| Methylpyruvate | - | - |
| Mono-Methylsuccinate | - | - |
| myo-Inositol; | + | + |
| N-Acetyl-D-Galactosamine | - | - |
| N-Acetyl-D-Glucosamine | + | + |
| N-Acetyl-D-mannosamine 6-phosphate | - | - |
| N-Acetyl-L-glutamate | - | - |
| N-Acetylneuraminate | - | - |
| Octopamine | - | - |
| Oxalate | - | - |
| Pectin | - | - |
| Phenylethylamine | - | - |
| Propane-1,2,3-tricarboxylate | - | - |
| Propane-1,2-diol | - | - |
| Propanoate | - | - |
| Putrescine | - | - |
| Pyruvate | - | - |
| Quinate | - | - |
| Ribitol | - | + |
| Salicin | - | + |
| Sebacic acid | - | - |
| Sedoheptulose | - | - |
| sn-Glycerol 3-phosphate | - | - |
| Stachyose | + | + |
| Starch | - | - |
| Succinate | - | - |
| Sucrose | + | + |
| Thymidine | - | - |
| Tyramine | - | - |
| Uridine | - | - |
| Xylitol | - | - |
| (R)-2-Methylmalate | - | - |
| (R)-Acetoin | - | - |
| (R,R)-Butane-2,3-diol | - | - |
| (R,R)-Tartaric acid | - | - |
| (S)-Lactate | - | - |
| 2-Amino-2-deoxy-D-gluconate | - | - |
| 2-Hydroxybenzoic acid | - | - |
| 2-Hydroxybutanoic acid | - | - |
| 2-Methylmaleate | - | - |
| 2-Oxobutanoate | - | - |
| 2-Oxoglutarate | - | - |
| 3-Hydroxyphenylacetate | - | - |
| 3-Oxalomalate | - | - |
| 4-Aminobutanoate | + | + |
| 4-Hydroxybenzoic acid | - | - |
| 4-Hydroxyphenylacetate | - | - |
| 5-Aminopentanoate | - | - |
| 5-Oxoproline | - | - |
| 6-Deoxy-D-galactose | - | - |
| **Nitrogen source** | ***In silico*** | **Experimental** |
| Ammonia | + | + |
| Nitrite | - | - |
| Nitrate | - | - |
| Urea | + | + |
| Biuret | - | - |
| L-Alanine | + | + |
| L-Arginine | + | + |
| L-Asparagine | - | + |
| L-Aspartic acid | + | + |
| L-Cysteine | - | - |
| L-glutamate | + | + |
| L-Glutamine | + | + |
| Glycine | + | + |
| L-Histidine | - | - |
| L-Isoleucine | - | + |
| L-Leucine | - | + |
| L-Lysine | - | + |
| L-Methionine | - | + |
| L-Phenylalanine | - | + |
| L-Proline | + | + |
| L-Serine | + | + |
| L-Threonine | + | + |
| L-Tryptophan | + | + |
| L-Tyrosine | - | + |
| L-Valine | - | + |
| D-Alanine | - | - |
| D-Aspartic acid | + | + |
| D-Glutamic acid | - | - |
| D-Lysine | - | - |
| D-Serine | + | + |
| D-Valine | - | - |
| L-Citrulline | + | + |
| L-Homoserine | + | + |
| L-Ornithine | + | + |
| N-Acetyl-L-Glutamic acid | + | + |
| N-Acetyl-L-glutamate | - | - |
| Hydroxylamine | - | - |
| Methylamine | - | - |
| Ethylamine | - | - |
| Ethanolamine | + | + |
| Putrescine | - | - |
| Agmatine | + | + |
| Histamine | - | - |
| Phenethylamine | - | - |
| Tyramine | - | - |
| Acetamide | - | - |
| Formamide | - | - |
| D-Glucosamine | + | - |
| N-Acetyl-D-Glucosamine | + | + |
| N-Acetyl-D-Galactosamine | - | - |
| N-Acetyl-D-Mannosamine | - | + |
| Adenine | - | - |
| Adenosine | - | - |
| Cytidine | - | - |
| Cytosine | - | - |
| Guanine | + | + |
| Guanosine | - | - |
| Thymine | - | - |
| Thymidine | - | - |
| Uracil | - | - |
| Uridine | - | - |
| Inosine | - | - |
| Xanthine | - | - |
| Xanthosine | - | - |
| Uric acid | + | + |
| Allantoin | + | + |
| 4-Aminobutanoate | + | + |
